# Music facilitates sleep initiation in adults with sleep-onset insomnia: The role of neural synchronization

**DOI:** 10.64898/2026.06.09.731083

**Authors:** Kira Vibe Jespersen, Alexandre Celma-Miralles, Peter Vuust

**Affiliations:** Center for Music in the Brain, Department of Clinical Medicine, Aarhus University and The Royal Academy of Music, Aarhus/Aalborg, Denmark

**Keywords:** Sleep, Music, Insomnia, Sleep onset, Neural synchronization, EEG, Beat perception

## Abstract

Sleep-onset insomnia is widespread in modern society, and many individuals turn to music to improve their sleep. While clinical studies have shown that music can positively affect sleep quality, the impact on sleep initiation remains unclear. Furthermore, there is limited knowledge on the mechanisms by which music may facilitate sleep. Here, we investigated whether music can facilitate sleep onset and if the effect is related to brain waves synchronizing to the slow beat of the sleep music. We recorded participants with sleep-onset insomnia (N=53) during a 30-minute afternoon rest using electroencephalography (EEG). Participants were randomly divided into two groups. Twenty-four participants listened to music chosen from a sleep music playlist while resting, and 29 rested in silence. We evaluated the transition from wakefulness towards sleep with the delta-alpha ratio of the EEG. To assess neural synchronization to the beat of the music, we used an EEG frequency tagging approach. We found a higher degree of transition towards sleep in the music group compared to silence over the 30-minutes resting period. Furthermore, higher beat stability in the music was reflected in stronger neural frequency tagging at the musical beat. However, the analyses showed no relationship between sleep initiation and neural synchronization to the beat. In sum, our results revealed that music has a positive effect on sleep initiation and that there is neural synchronization to naturalistic sleep music with a steady beat, but we found no indication that this neural synchronization is the central mechanism driving enhanced sleep initiation with music.

## 1 Introduction

Sleep-onset insomnia is a prevalent and increasing problem in modern society, highlighting the need for safe and accessible interventions (Garland et al., 2018; Jensen, Davidsen, Christensen, Ekholm, & Christensen, 2026; Pallesen, Sivertsen, Nordhus, & Bjorvatn, 2014; Sivertsen, Harvey, Vedaa, Pallesen, & Hysing, 2026). Listening to music is widely used as a strategy to facilitate sleep (Buus, Genovese, & Jespersen, 2025; Morin, LeBlanc, Daley, Gregoire, & Mérette, 2006), and a growing body of research suggests that music may improve sleep quality across a range of populations, including individuals with insomnia, older adults, hospitalized patients, and individuals with mental health disorders (C. T. Chen et al., 2021; Jespersen, Hansen, & Vuust, 2023; Jespersen, Pando-Naude, Koenig, Jennum, & Vuust, 2022; Kakar, Venema, Jeekel, Klimek, & van der Jagt, 2021; Zhao, Lund, & Jespersen, 2024). However, despite this growing interest, the extent to which music specifically facilitates the transition from wakefulness to sleep remains unclear.

While subjective reports suggest that music can help individuals fall asleep faster (Trahan, Durrant, Müllensiefen, & Williamson, 2018), objective evidence is more limited and inconclusive. A small number of studies using electroencephalography (EEG) have examined the effects of music on sleep onset latency, yielding largely mixed or null results in healthy young adults (Chang, Lai, Chen, Hsieh, & Lee, 2012; C. K. Chen et al., 2014; Cordi, Ackermann, & Rasch, 2019; Costa et al., 2025; Lazic & Ogilvie, 2007). Similarly, intervention studies including clinical populations have reported inconsistent findings (Jespersen, Otto, Kringelbach, Van Someren, & Vuust, 2019; Su et al., 2013), and none have focused specifically on sleep-onset insomnia. As such, the effectiveness of music as a tool for promoting sleep initiation remains insufficiently established, particularly in clinically relevant populations.

Beyond the question of efficacy, the neurophysiological mechanisms by which music may influence sleep initiation are poorly understood. One proposed mechanism is neural synchronization to the beat of the music, whereby ongoing brain oscillations align with the temporal regularities of the auditory input (Mayayo, Celma-Miralles, Keller, & Toro, 2026; Møller et al., 2026). This alignment leads to enhanced neural activity at beat-related frequencies (Celma-Miralles, Seeberg, Haumann, Vuust, & Petersen, 2024; Nozaradan, 2012; Sifuentes-Ortega, Lenc, Nozaradan, & Peigneux, 2022). This process, often referred to as neural entrainment, is a well-established phenomenon in music perception(Celma-Miralles & Toro, 2019; Nozaradan, 2012, 2014; Sauvé, Bolt, Nozaradan, & Zendel, 2022) and has been suggested as a potential pathway through which music could facilitate sleep onset (Dickson & Schubert, 2019; Jespersen, 2022; Lazic & Ogilvie, 2007). In the context of sleep, it has been hypothesized that slow musical rhythms may promote sleep by enhancing low-frequency neural activity characteristic of the sleeping brain. However, empirical evidence directly linking beat-related neural synchronization to sleep initiation is scarce, and existing studies have yielded mixed findings (Costa et al., 2025; Lazic & Ogilvie, 2007).

All together this shows that although music is commonly reported to facilitate sleep, objective evidence for its effects on sleep initiation is limited and inconsistent. Existing EEG studies have produced mixed results and have focused on healthy participants rather than individuals with sleep-onset insomnia. As a result, it remains unclear whether music effectively promotes sleep initiation in this clinical population and through which mechanisms such effects might arise.

In the present study, we investigated whether listening to music facilitates the transition from wakefulness to sleep in adults with sleep-onset insomnia, and whether neural synchronization to the musical beat constitutes a potential mechanism underlying this effect. Using EEG-based measures, we tested the hypotheses that (1) music promotes sleep initiation compared to silence, and (2) neural synchronization to the beat underlies the sleep promoting effect of music.

## 2 Methods

### 2.1 Participants

The study included EEG recordings from a total of 53 adult participants with sleep-onset insomnia. Participants were recruited via social media announcements. To be included participants had to be between 18 and 65 years old and fulfil the DSM-5 diagnostic criteria for insomnia disorder with sleep initiation difficulties and symptom duration of max 18 months. Exclusion criteria were use of medications affecting sleep, hearing problems, abuse of alcohol or drugs, somatic disorders interfering with sleep, current psychiatric disorders, shift work, pregnancy and other sleep disorders. All participants underwent a clinical interview as well one-night ambulant polysomnography screening for breathing- and motor-related sleep disorders. The included participants were between 19 and 55 years old with a majority of females (see Table 1).

**Table 1.** Participant characteristics.

| | All<br>(N = 53) | Music<br>(n = 24) | Control<br>(n = 29) | Test $p$-value |
| --- | --- | --- | --- | --- |
| Age | 28.87 (8.65) | 27.83 (6.50) | 29.72 (10.12) | $W = 340$<br>$p = 0.893$ |
| Sex, male/female | 16/37 | 6/18 | 10/19 | $\chi^2 = 0.20$<br>$p = 0.554$ |
| Insomnia severity* | 15.36 (4.20) | 15.42 (3.91) | 15.31 (4.49) | $t = 0.09$<br>$p = 0.928$ |
| Musicianship** (n) |  |  |  |  |
| -Nonmusician | 7 | 5 | 2 |  |
| -Music-loving nonmusician | 32 | 14 | 18 |  |
| -Amateur musician | 10 | 4 | 6 | $\chi^2 = 3.07$ |
| -Serious amateur musician | 3 | 1 | 2 | $p = 0.579$ |
| -Semiprofessional musician | 1 | 0 | 1 |  |
| -Professional musician | 0 | 0 | 0 |  |

Data is reported as mean and standard deviation (SD) unless otherwise specified.
\*Measured with the Insomnia Severity Index (Bastien, Vallieres, & Morin, 2001). Higher scores indicate more severe insomnia.
\*\*Item from the Ollen Musical Sophistication Index (Ollen, 2006).

Participants were randomized to two different conditions. The music group included 24 participants, whereas 29 participants were in the control group resting in silence. In the music group, 12 participants chose the Minimalistic sleep playlist and 12 participants chose the New Age playlist (see Table S1 for music specifications). Of the 53 participants, 12 participants had missing data at one timepoint due to technical issues (n = 4) or because they missed their appointment (n = 8). Reasons for missing the appointment included sickness and covid isolation (n = 4) and dropout (n = 4).

As part of a larger randomized controlled trial, the study was approved by the Ethical Committee of the Central Denmark Region (ID 1-10-72-32-20). In accordance with the Helsinki Declaration, all participants received oral and written information about the study and signed informed consent before enrolment.

### 2.2 Procedure

Upon enrolment participants were randomized to either music or control group using permuted block randomization with variable block sizes implemented in REDCap (Harris et al., 2019). Participants in the music group were presented with a choice among 5 playlists of different genres and could choose the one they liked the best. The music in all playlists was compiled to facilitate arousal reduction based on the criteria of a slow tempo, stable dynamics, simple structure, no text or lyrics (Bernardi et al., 2009; Bernardi, Porta, & Sleight, 2006; Gomez & Danuser, 2007; Trehub & Trainor, 1998).

We measured EEG activity during a 30-minute afternoon rest. Participants were placed comfortably lying down in supine position in a bed in a separate room with room temperature controlled at a stable level of 22 degrees. Participants were instructed to relax with eyes closed. They were given no instructions to sleep, but if they asked, they were told that they could sleep or not as they preferred. All recordings were done between 1 pm and 5 pm. Participants in the music group listened to their selected music during the 30-minutes rest, and participants in the control group rested in silence.

Music stimuli were presented with PsychoPy3 v2020.1.1 using etymotic in-ear headphones. Because the study was embedded in an RCT trial, the recordings were repeated at two sessions, one at baseline and one four weeks later with the same procedure. During the four weeks between recordings, participants in the music group had been instructed to listen to their selected playlist every night at bedtime.

### 2.3 EEG recordings

Continuous EEG was recorded using a 32-channel Acticap system from Brainvision with a 1000 Hz sampling rate. Channels were positioned and labelled according to the 10/20 system. Reference was FCz and the montage included vertical and horizontal electrooculogram channels.

EEG preprocessing consisted of filtering the data between 0.1 and 45 Hz (filter order = 4). A 50 Hz and harmonics notch filter was applied to reduce line noise, bad channels were removed and interpolated and independent component analysis (ICA) decomposition was applied under the “fastica” option. Components capturing eye movements or single-channel artefacts were removed from the data by visual inspection of the components’ timeseries and topographies. Afterwards, channels were re-referenced to the common average and mastoids and EOG were removed from the dataset.

For group comparison analyses, the EEG recordings from both music and control groups were divided into ten consecutive 3-minute segments. For steady state evoked potential (SSEP) analyses linking neural amplitudes with the beat of music, the EEG recordings from the music group of participants were segmented in accordance with the length of each music track, resulting in ten tracks lasting approximately 3 minutes each. EEG preprocessing and analyses were done using custom Matlab scripts (The MathWorks, Inc., Natick, Massachusetts, US).

### 2.4 Power spectral density

We applied power spectrum density analyses to the preprocessed EEG epochs to measure the activity of the 30 electrodes at 5 frequency bands: delta (1-4 Hz), theta (4-8 Hz), alpha (8-12 Hz), beta (12-30 Hz) and gamma (30-45 Hz). We computed the relative power of each frequency band by dividing its activity over the total activity across the 5 frequency bands (i.e., between 1 and 45 Hz). We then converted the measures into percentages by multiplying them by 100 (Comsa, Bekinschtein, & Chennu, 2019; Šušmáková & Krakovská, 2007). Before averaging electrodes at each frequency band and epoch, we removed those that showed power with 1.5 interquartile ranges above the upper quartile (75 percent) or below the lower quartile (25 percent) using the Matlab function *isoutlier*(data,’quartiles’). The resulting relative power of all frequency bands is depicted in Figure S1 for each group of participants.

### 2.5 Wake/sleep transition

To have a single sensitive measure evaluating the level of wakefulness and potential transition towards sleep during the 30-minutes rest period, we computed the ratio between delta and alpha relative power (Prerau et al., 2014; Šušmáková & Krakovská, 2007). Most people show enhanced alpha activity when relaxing with eyes closed (Berger, 1929; Hohaia, Saurels, Johnston, Yarrow, & Arnold, 2022). However, sleep onset is characterised by a decrease in alpha power and an increase of slow-wave activity (Lacaux, Strauss, Bekinschtein, & Oudiette, 2024). Therefore, this ratio approach – also termed the alpha slow-wave index - is commonly used in research evaluating the sleep onset process (Abbattista, Wacquier, & Strauss, 2026; Comsa et al., 2019; Jobert et al., 1994; Li et al., 2025; Prerau et al., 2014; Strauss, Sitt, Naccache, & Raimondo, 2022). In addition, sleep scoring of the EEG data was done using the Usleep online platform v2.0 developed at University of Copenhagen (https://sleep.ai.ku.dk) (Perslev et al., 2021). The automatic sleep scoring was done in 30-second epochs based on channels Fz, F3, F4, Cz, C3, C4, Pz, P3, P4, Oz, O1, O2 and EOG recordings.

### 2.6 Steady State Evoked Potentials

For the EEG segments corresponding to the music tracks, we removed the first 2 seconds of each trial to discard the evoked potentials related to track change. For each participant and electrode, we applied a fast Fourier transform with a zero-padding of 300000 to homogenize all the segments up to 5 minutes. This transformed the amplitudes over time (μV/s) into amplitudes over frequencies (μV/Hz), with a frequency resolution of 0.0033 Hz. According to the frequency tagging approach (Nozaradan, 2014; Sifuentes-Ortega et al., 2022), these amplitudes reflect stimulus-related neural activity mixed with spontaneous unrelated activity and background noise. We normalized the output of the frequency transformation across frequencies by subtracting the mean value of non-adjacent (between 4 and 9) surrounding frequency bins at each frequency bin. That is, the mean of the amplitudes falling between -29.7 and - 13.2 mHz and 13.2 and 29.7 mHz (milliHertz). This increases the signal-to-noise ratio, because if no steady-state evoked potentials are present, the voltage amplitudes may vary similarly across frequencies after the subtraction (i.e., amplitudes around zero), but if some periodicities are consistent in the auditory signal, their corresponding peaks should remain after the subtraction (i.e., amplitudes greater than zero). See Figure 2 for a visualization of the approach with an example of peaks related to the beat of a track in the auditory stimulus and the corresponding EEG. To account for any spectral leakage, we calculated the mean of the amplitudes of 3 frequency bins centered at each frequency of interest, summing those for each assumed beat and its first harmonic (Retter, Rossion, & Schiltz, 2021), and averaged them across eight frontocentral channels (F3, Fz, F4, FC1, FC2, C3, Cz, C4). Previous work supports that beat-related activity mainly occurs on this topographical location (Celma-Miralles et al., 2024; Nozaradan, Peretz, & Mouraux, 2012; Sauvé et al., 2022; Stupacher, Witte, Hove, & Wood, 2016). Finally, to confirm the presence of periodicities related to the beat of music, we selected the beat-related amplitudes for each track and tested whether these amplitudes were greater than zero using a linear mixed-effect model with the intercept centered at 0:

*model_beatAmp <- lmer(beatAmplitude ∼ 0 + music track + (session* | *participant)*

**Figure 1.**
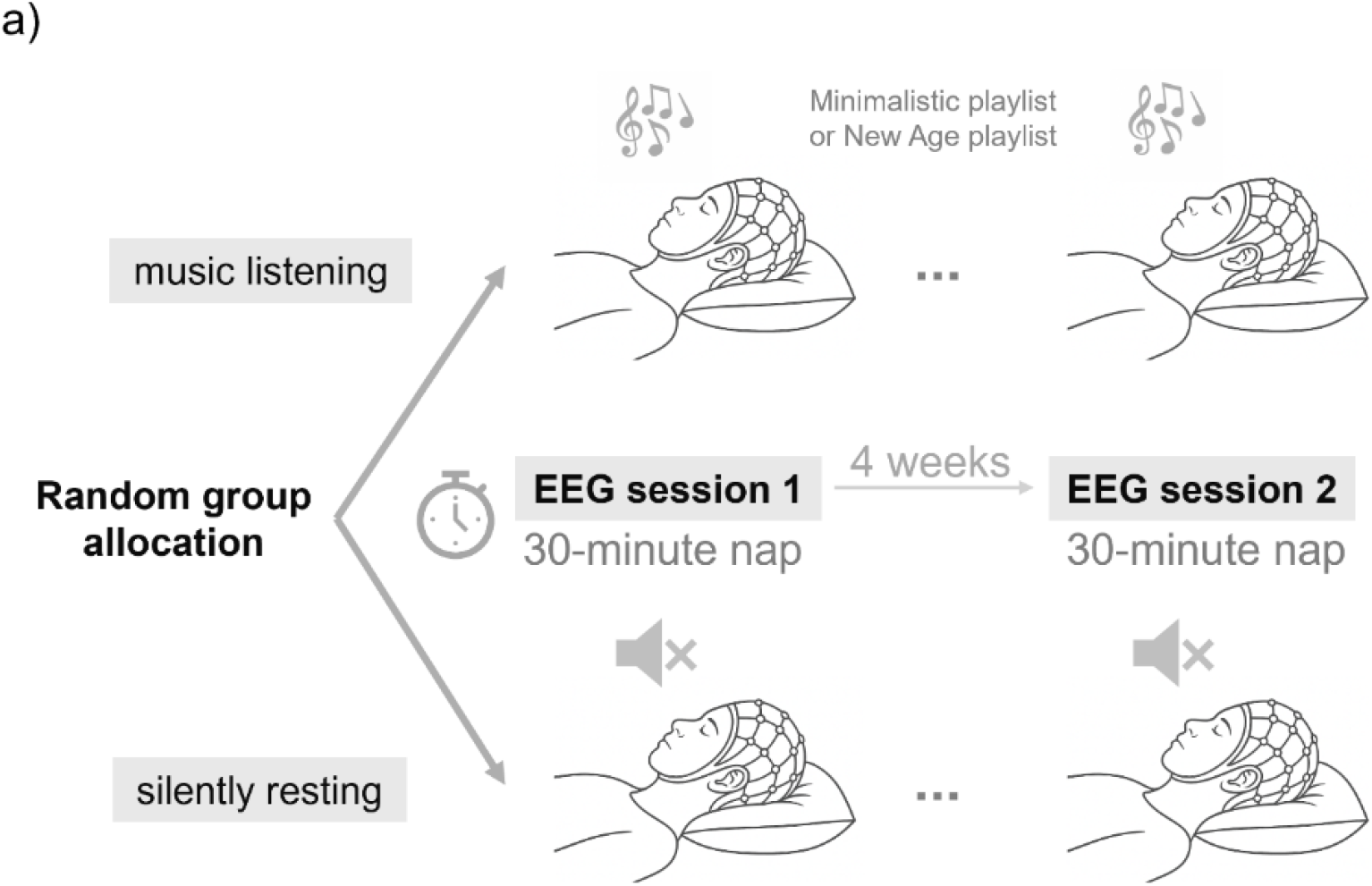
Study design. (a) Participants were asked to rest in the lab, either listening to music or in silence, while their EEG was recorded for 30 minutes in two separated sessions.

**Figure 2.**
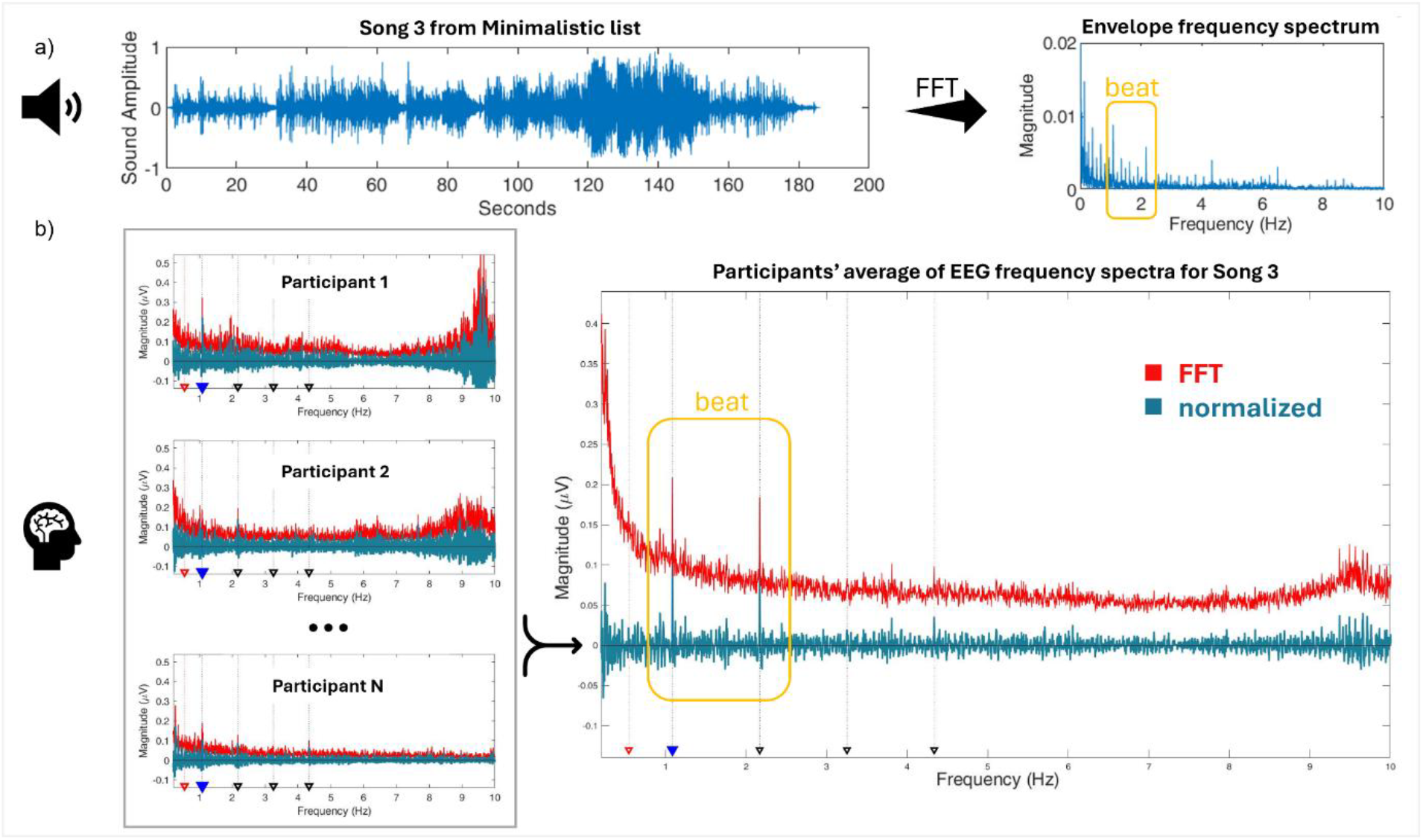
Beat-related amplitudes in an example music track and its corresponding EEG activity. (a) The envelope of the sound waveform is converted into a frequency spectrum to identify the peaks related to beat periodicities. (b) EEG recordings of different individuals and their mean when they are listening to the same music track. Neural responses are frequency-tagged to measure the amplitudes related to the identified beat periodicities: the blue triangle represents the expected beat and its adjacent black triangle represents its harmonic. Results from the fast Fourier transform are in red, and results after applying the neighbour-subtraction as signal-to-noise normalization are in turquoise.

### 2.7 Beat stability

To address tempo fluctuations across the music playlists, the consistency of the beat in each music track was determined by sensorimotor synchronization (i.e., finger tapping responses) based on the two periodicities obtained from the audio spectrograms (see Supplementary Figure S2 and Figure 4c). Beat stability of each track was defined with circular analyses of the tapping responses of 10 pilot participants. The consistency of the taps was operationalized as the mean vector length (i.e., 1- variance) after all the taps were transposed on a circle in relation to their “time” of occurrence within the periodicity of the beat and its double periodicity. The timings of the taps were converted into angular measures (i.e., phase in radians) by calculating the modulus of each tap in reference to the tempo in milliseconds (i.e., the inter-onset interval of the beat or its double periodicity (Møller, Stupacher, Celma-Miralles, & Vuust, 2021). The length of the vectors represents the consistency of the taps at each periodicity (Figure S2 and Table S1).

### 2.8 Statistical evaluation

Statistical analyses were done in R (Rstudio 2026.01.0 build 392, Posit Software PBC). For the analysis of the effect of music on the sleep onset process, we created a mixed-effects model with Group, Time and Session as fixed effects and random effects for Participants and Segments and random slope for Session (see Table S2). To evaluate the impact of music on the level of wakefulness (delta-alpha ratio) during the 30-minute rest, we looked at Group x Time interaction. The Time dimension reflected the progression over the ten 3-minutes segments and was modelled both with a linear effect and a quadratic effect. Model fit was evaluated using AIC and BIC.

We applied repeated measures correlations between the log-transformed delta-alpha ratio and beat-related neural amplitudes for each track using the R package *rmcorr* (Bakdash & Marusich, 2017). To see if neural synchronization to the beat facilitated transitioning towards sleep, we tested if the delta-alpha ratio, i.e. the degree of wakefulness/sleep, of a segment correlated with the beat-related neural amplitude of the previous track. Finally, we tested correlations between beat-related neural responses for each music track and the beat stability.

## 3 Results

### 3.1 Transition from wakefulness towards sleep with and without music

To test the first hypothesis, we used a linear mixed effect model evaluating the evolution of the delta-alpha ratio over the 30-minute rest time in both groups. The model showed a main effect of Time reflecting a general transition towards sleep indicated by a higher delta-alpha ratio, i.e. an increase in delta activity and/or a reduction in alpha activity (*β* = 6.018, *t* = 4.70, *p* < 0.001. There was no main effect of Group, but the Group x Time interaction was significant (*β* = 2.976, *t* = 2.38, *p* = 0.017). This reflects a higher degree of sleep initiation over time in the music group compared to participants resting in silence (Figure 3). There was no significant effect of session, meaning that this pattern was the same before and after daily exposure to the music in the music group (Table S2). Analyses of the percentage of time spent asleep based on sleep scores showed the same results (Figure S3 and Table S3). We did not see any significant differences between groups when comparing power spectral density of each frequency band (Tables S4-S8).

**Figure 3.**
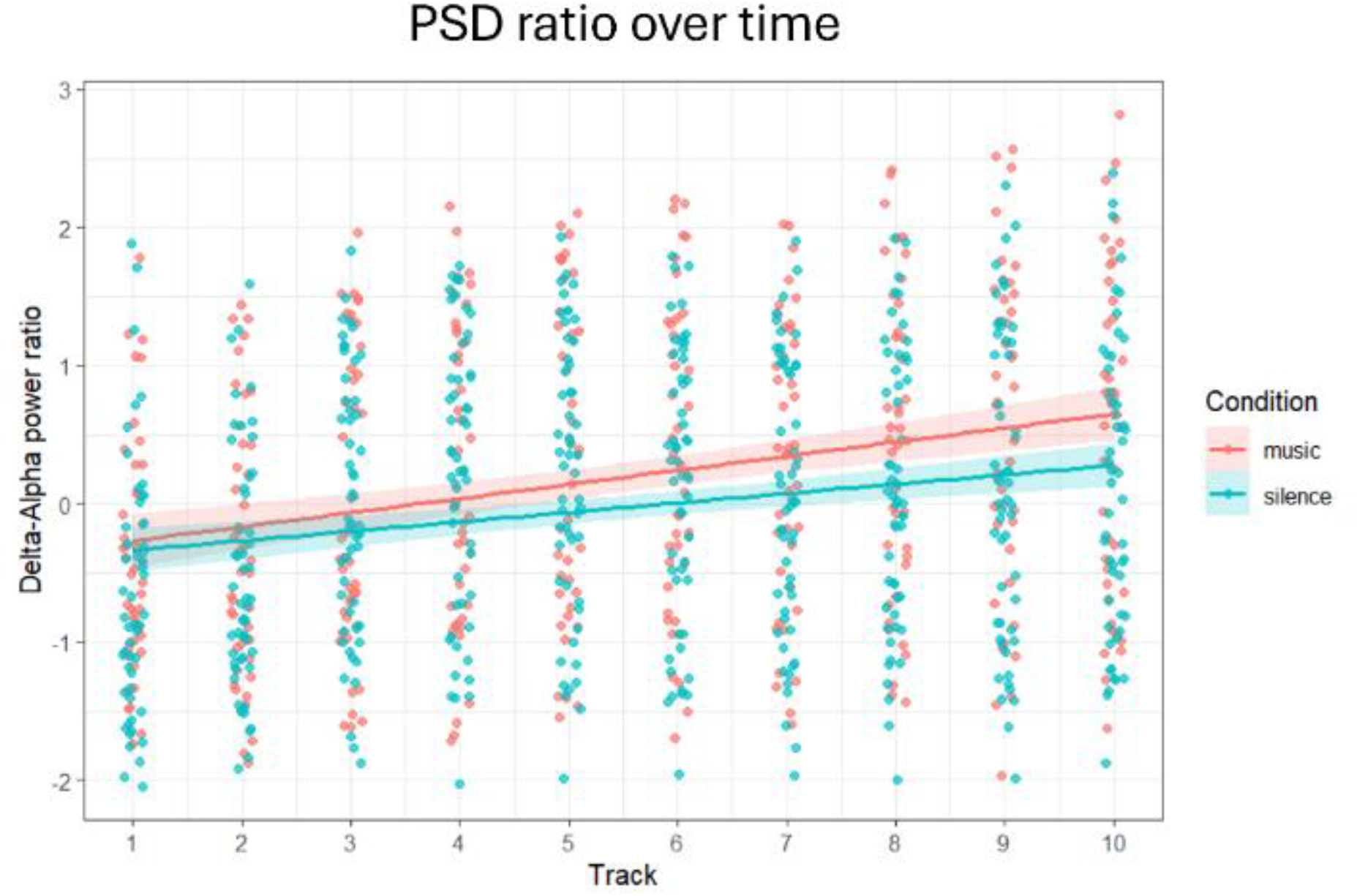
Delta-Alpha ratio over time in each group. The music group (salmon) has a higher increase in Delta-Alpha ratio compared to the silence group (turquoise) (β = 2.976, t = 2.38, p = 0.017). Higher Delta-alpha ratio reflects sleep initiation. As such, these findings show that participants listening to music transition more towards sleep compared to those resting in silence. The 30-minute recording is segmented into 10 segments each 3 minutes long.

### 3.2 Is the sleep promoting effect of music driven by neural synchronization to the beat?

To investigate whether the sleep facilitating effect of music was related to neural synchronization to the beat, we focused on the participants in the music group using a frequency tagging approach (Figure 2). For all music tracks of the two sleep playlists selected by participants, the beat-related frequencies (beat and double beat) were in the delta range (Table S1). The frequency tagging results showed enhanced beat-related amplitude, reflecting neural synchronization to the beat, in some tracks, but not all. In the Minimalistic playlist there was enhanced beat-related amplitude only in track 3 (t = 9.37, p < 0.001). In the New Age playlist, we found enhanced beat-related amplitudes for tracks 1, 2, 3, 4, 5, 6, 7 and 10 (Table S1 and Table S9).

We used repeated measures correlations to test whether there was any relationship between the neural synchronization to the beat and the following delta-alpha ratio indicating transitioning towards sleep. The results showed no correlation between the two measures (r (327) = -0.02, 95% CI: -0.13 to 0.08, *p* = 0.678) and hence did not support the hypothesis of neural synchronization underlying the impact of music on sleep onset (Figure S4).

### 3.2.1 Is the effect occluded by sleep onset?

To explore alternative explanations for our results, we investigated if the results were affected by the influence of sleep onset on neural synchronization. Previous research has found that neural synchronization to beat-related frequencies is attenuated with sleep onset (Sifuentes-Ortega et al., 2022). We therefore tested if the neural synchronization to the beat only in tracks characterised as wakefulness predicted delta-alpha ratio and thereby transitioning towards sleep in the following track. However, the results still did not show any relationship between neural synchronization to beat-related frequencies and sleep initiation (r(290) = -0.0001, 95% CI: -0.13 to 0.13, p = 0.999).

To see if our data aligned with the findings of Sifuentes-Ortega and colleagues, we evaluated the relationship between the delta-alpha ratio and the degree of neural synchronization. Here, we found a trend towards a negative correlation between the delta-alpha ratio and beat-related amplitudes (*r*(385) = -0.095, *p* = 0.063, Figure 4d). This indicates reduced neural synchronization to the beat when transitioning from full wakefulness towards sleep, similar to the results of Sifuentes-Ortega and colleagues.

**Figure 4.**
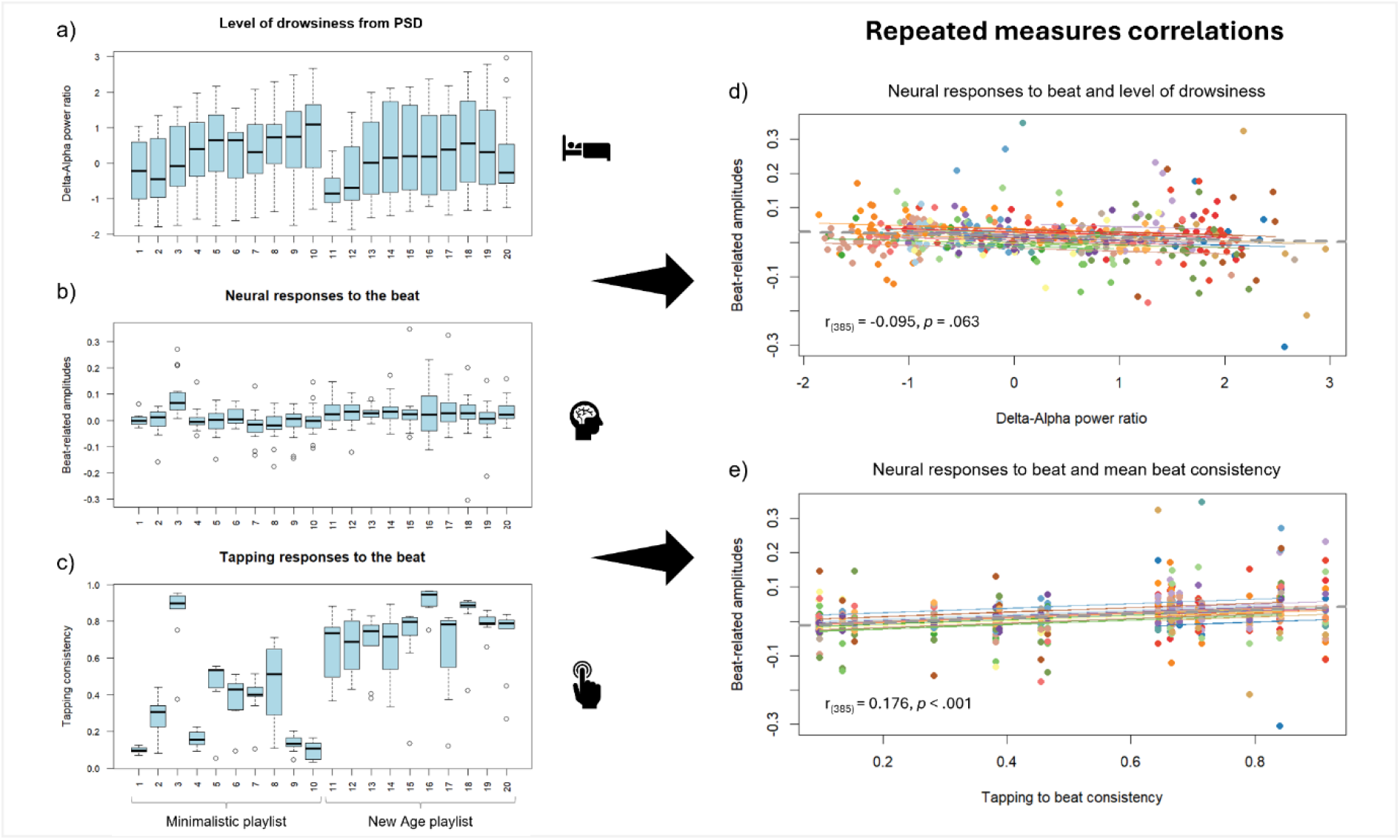
Repeated measures correlations between beat-related amplitudes and delta-alpha ratio or beat consistency at each song of the Minimalistic and New Age playlists. In the boxplots, the x-axis reflects each track of the two playlists starting with the Minimalistic playlist (1-10) and followed by the New Age playlist (11-20). Each track is approximately 3 minutes long. (a) The delta-alpha ratio (log-transformed) of the participants (y-axis) increases over time both in participants listening to the Minimalistic playlist (x-axis: 1-10) and those listening to the New Age playlist (x-axis: 11-20). In the New Age group, participants seem to be returning towards more wakefulness during the last track. (b) The beat-related amplitudes (y-axis) for each track show differing degrees of neural synchronization to the beat. Most tracks in the New Age playlist (x-axis: 11-20) are significantly enhanced, whereas only track 3 in the Minimalistic playlist is significantly different from 0 (Table S9). (c) We evaluated beat stability by having a control group of 10 participants tap along to each track. Tapping consistency for each track reflects beat stability. (d) Repeated measures correlations between beat-related amplitudes on the y-axis and the logarithmically transformed delta-alpha ratio on the x-axis. The results show a trend towards a negative correlation potentially indicating reduced neural synchronization to the beat when transitioning towards sleep. (e) When looking at the relationship between neural synchronization (y-axis) and beat stability (x-axis), we see a significant correlation. The more stable the music beat, the stronger the neural activity at beat-related frequencies.

#### 3.2.2 Does beat stability play a role?

Since participants listened to naturalistic music, another potentially relevant factor is beat stability. Previous studies using frequency tagging approaches have focused on isochronous beats and stable rhythms designed for the specific study (Lenc, Keller, Varlet, & Nozaradan, 2020; Møller et al., 2026; Nozaradan, 2012; Sifuentes-Ortega et al., 2022). Here, we used naturalistic music selected for sleep, and above we saw that there was neural synchronization only to some of the sleep music tracks.

We evaluated beat stability using finger tapping and circular statistics (Figure S2 and Figure 4C). In a repeated measures correlation analysis, we found a significant correlation between beat stability and the amplitude of the beat-related amplitudes in the EEG (r = 0.176, p < 0.001; Figure 4E). This result reflects a higher degree of neural synchronization to the beat in tracks with a stable beat compared to music tracks with a less stable beat, e.g. music with rubato, ritardando and accelerando.

## 4 Discussion

We found that listening to sleep music promoted the transitioning from wakefulness to sleep in adults with sleep-onset insomnia. Compared to the group resting in silence, the music-listening group showed a greater increase in the delta-alpha ratio reflecting sleep initiation. We found no evidence supporting the hypothesis that neural synchronization to the music beat may facilitate sleep initiation. Instead, the synchronized neural activity rather reflected the stability of the beat in the tracks of the naturalistic sleep music playlists.

The finding that music promotes the transitioning towards sleep over time suggests that music can be useful as an intervention facilitating sleep initiation. This is in line with subjective reports showing that many people who use music as a sleep aid experience that the music helps them fall asleep faster (Trahan et al., 2018). However, we did not see a main effect of music intervention which is consistent with previous studies showing mixed results of the effect of music on general measures such as sleep onset latency (Cordi et al., 2019; Costa et al., 2025; Jespersen et al., 2019; Lazic & Ogilvie, 2007). Instead, the delta/alpha ratio increased more with music than without it during the course of the rest period. Similarly, the percentage of time scored as sleep increased more in the music group compared to the participants resting in silence. This highlights the need for analyses taking the time dimension into account in order to clarify the effect of music on sleep initiation. Here, we used adult participants with clearly defined sleep-onset insomnia. Therefore, our results indicate that music can impact the transitioning from wakefulness to sleep in this population and highlights the relevance of these findings in a clinical context.

Contrary to our expectations, we did not find any evidence that neural synchronization to the beat plays a direct role in the somnogenic effect of music. Therefore, our results do not support the hypothesis that neural synchronization to the beat is a central mechanism underlying the ability of music to promote sleep initiation. Existing evidence linking beat-related neural activity to sleep onset is limited. By applying frequency-tagging analyses to link neural activity at specific beat-related frequencies –rather than broader delta-band activity in general–, our study contributes to the field by using a novel but well-established methodology to investigate the neurophysiological mechanisms underlying the impact of music on sleep.

Previous studies have applied a broader approach testing the ability of music to promote EEG synchronization in the delta or sub-delta range (0.2 Hz - 3.5 Hz) (Costa et al., 2025; Lazic & Ogilvie, 2007). These studies showed that listening to slow music could enhance delta-band activity to some degree. However, in our study we did not find generally enhanced delta activity in the music group compared to the silence group. Our PSD analyses showed a general increase in delta activity over time, and just a trend towards an interaction effect with a higher increase in delta activity in the music group (Table S4). Several differences between our study and previous studies may explain the differing results regarding delta activity. First, we recorded EEG during a 30-minutes afternoon rest and not a full night of sleep. Secondly, we included a sample of adults with sleep-onset insomnia instead of healthy young adults and used a between-group randomized design rather than a within-subject design. The within-subject design used in the other two studies can reduce variability and may thereby strengthen statistical power to detect differences. However, we included more participants (N=53) than in the other two studies (N= 10 and N=22, respectively). Finally, the previous studies used a standard polysomnography approach including six or eight EEG electrodes, and the delta frequency results were significant only in one channel. In contrast, we used a full 32- channel EEG cap.

Our results showed varying degrees of neural synchronization to the beat but found no indication that these synchronization measures were related to the sleep promoting effect of the music. This difference in the degree of neural synchronization could stem from the interplay of two relevant factors: beat stability (i.e., greater tempo fluctuations in the Minimalistic than in the New Age playlist) and weakened synchronized neural activity due to the sleep states of the participants (Sifuentes-Ortega et al., 2022). Indeed, we found that when using naturalistic sleep music, stimuli neural synchronization was strongly related to beat stability of the presented track (Figure 4e). This is in line with previous studies showing stronger neural synchronization with more rhythmically stable stimuli (Kaneshiro, Nguyen, Norcia, Dmochowski, & Berger, 2020; Lenc et al., 2020). Similarly, we investigated if sleep onset affected the degree of neural synchronization to the beat. Previous research has shown that the well-documented enhancement of beat-related activity seen during wakefulness, is attenuated during sleep (Sifuentes-Ortega et al., 2022). They showed that neural amplitudes of syncopated and unsyncopated rhythms seen during wakefulness were weakened in the REM sleep stage and fully suppressed during non-REM sleep. Our results only showed a small trend towards lower neural amplitudes at beat-related frequencies with increased delta-alpha ratio, and as such, we do not fully replicate their findings. Again, our study presents crucial differences compared to the paradigm of Sifuentes-Ortega and colleagues. We focused on sleep initiation during a 30-minute afternoon rest in participants with sleep-onset insomnia, which substantially differs from their reported full night EEG recordings in participants with good sleep quality. Furthermore, the stimuli we used were naturalistic music, played on several instruments and including tempo changes between and within tracks, instead of repetitive lab-designed rhythmic patterns played on a single tone. Due to the 30-minute recording session, and the length of each music track, we could not perform frequency tagged comparisons of every single sleep stage from the sleep scores, and rather focused on the evolution of delta-alpha ratio as an indicator of transitioning towards sleep.

## Conclusion

Epidemiological evidence indicates a growing prevalence of insomnia, highlighting the need for safe and accessible interventions. In this study, we tested whether listening to music facilitates the transition from wakefulness to sleep in individuals with sleep-onset insomnia, and whether neural synchronization to the beat of the music constitutes a potential underlying mechanism. While neural synchronization to the beat was not associated with sleep initiation, listening to naturalistic sleep music promoted the transition from wakefulness to sleep as measured by EEG. Together with previous research, these findings suggest that music listening can support sleep initiation processes and represents a promising low-cost, non-pharmacological aid for individuals with sleep-onset insomnia.

## Acknowledgments

We thank members of MIB for valuable comments and students who helped with the EEG preprocessing.

## Conflicts of Interest

The authors declare no potential conflicts of interest with respect to the research, authorship, and/or publication of this article.

## Funding

Center for Music in the Brain was funded by the Danish National Research Foundation (DNRF117), the Lundbeck Foundation (R469-2024-1573), and Købmand Herman Sallings Fond.

## Data Availability Statement

The data that supports the findings of this study are available from the corresponding authors upon reasonable request.

## Notes

### Competing Interest Statement

The authors have declared no competing interest.

## References

Abbattista, L., Wacquier, B., & Strauss, M. (2026). Hypervigilance profiles in sleep-onset insomnia and psychiatric comorbidity. bioRxiv, 2026.2005. 2005.722943.

Bakdash, J. Z., & Marusich, L. R. (2017). Repeated measures correlation. Frontiers in Psychology, 8, 456.

Bastien, C. H., Vallieres, A., & Morin, C. M. (2001). Validation of the Insomnia Severity Index as an outcome measure for insomnia research. Sleep Medicine, 2(4), 297–307.

Berger, H. (1929). Über das elektroenkephalogramm des menschen. Archiv für Psychiatrie und Nervenkrankheiten, 87(1), 527–570.

Bernardi, L., Porta, C., Casucci, G., Balsamo, R., Bernardi, N. F., Fogari, R., & Sleight, P. (2009). Dynamic interactions between musical, cardiovascular, and cerebral rhythms in humans. Circulation, 119(25), 3171–3180. doi:10.1161/CIRCULATIONAHA.108.806174

Bernardi, L., Porta, C., & Sleight, P. (2006). Cardiovascular, cerebrovascular, and respiratory changes induced by different types of music in musicians and non-musicians: the importance of silence. Heart, 92(4), 445–452. doi:10.1136/hrt.2005.064600

Buus, R. M., Genovese, S., & Jespersen, K. V. (2025). The art of sleep: examining sleep strategies in the general population with a focus on the use of music for sleep. Journal of Sleep Research, e70006.

Celma-Miralles, A., Seeberg, A. B., Haumann, N. T., Vuust, P., & Petersen, B. (2024). Experience with the cochlear implant enhances the neural tracking of spectrotemporal patterns in the Alberti bass. Hearing Research, 452, 109105. doi:10.1016/j.heares.2024.109105

Celma-Miralles, A., & Toro, J. M. (2019). Ternary meter from spatial sounds: Differences in neural entrainment between musicians and non-musicians. Brain and Cognition, 136, 103594.

Chang, E. T., Lai, H. L., Chen, P. W., Hsieh, Y. M., & Lee, L. H. (2012). The effects of music on the sleep quality of adults with chronic insomnia using evidence from polysomnographic and self-reported analysis: A randomized control trial. International Journal of Nursing Studies, 49(8), 921–930. doi:10.1016/j.ijnurstu.2012.02.019

Chen, C. K., Pei, Y. C., Chen, N. H., Huang, L. T., Chou, S. W., Wu, K. P., … Wu, C. K. (2014). Sedative music facilitates deep sleep in young adults. Journal of Alternative and Complementary Medicine, 20(4), 312–317. doi:10.1089/acm.2012.0050

Chen, C. T., Tung, H. H., Fang, C. J., Wang, J. L., Ko, N. Y., Chang, Y. J., & Chen, Y. C. (2021). Effect of music therapy on improving sleep quality in older adults: A systematic review and meta-analysis. Journal of the American Geriatrics Society, 69(7), 1925–1932.

Comsa, I. M., Bekinschtein, T. A., & Chennu, S. (2019). Transient topographical dynamics of the electroencephalogram predict brain connectivity and behavioural responsiveness during drowsiness. Brain Topography, 32(2), 315–331.

Cordi, M. J., Ackermann, S., & Rasch, B. (2019). Effects of relaxing music on healthy sleep. Scientific Reports, 9(1), 1–9.

Costa, M., Andreose, A., Barzetta, F., Beracci, A., Fabbri, M., Ferri, R., … Martoni, M. (2025). Slow pentatonic sequences facilitate sleep onset. Psychology of Music, 03057356251359079.

Dickson, G. T., & Schubert, E. (2019). How Does Music Aid Sleep?,Literature Review. Sleep Medicine.

Garland, S. N., Rowe, H., Repa, L. M., Fowler, K., Zhou, E. S., & Grandner, M. A. (2018). A decade’s difference: 10-year change in insomnia symptom prevalence in Canada depends on sociodemographics and health status. Sleep health, 4(2), 160–165.

Gomez, P., & Danuser, B. (2007). Relationships between musical structure and psychophysiological measures of emotion. Emotion, 7(2), 377–387. doi:10.1037/1528-3542.7.2.377

Harris, P. A., Taylor, R., Minor, B. L., Elliott, V., Fernandez, M., O’Neal, L., … Kirby, J. (2019). The REDCap consortium: building an international community of software platform partners. Journal of Biomedical Informatics, 95, 103208.

Hohaia, W., Saurels, B. W., Johnston, A., Yarrow, K., & Arnold, D. H. (2022). Occipital alpha-band brain waves when the eyes are closed are shaped by ongoing visual processes. Scientific Reports, 12(1), 1194.

Jensen, H. A. R., Davidsen, M., Christensen, A. V., Ekholm, O., & Christensen, A. I. (2026). Danskernes Sundhed – Den Nationale Sundhedsprofil 2025. Retrieved from https://www.sst.dk/sundhedsprofilen

Jespersen, K. V. (2022). A Lullaby to the Brain: The Use of Music as a Sleep Aid. In B. Colombo (Ed.), The Musical Neurons (pp. 53–63). Cham: Springer International Publishing.

Jespersen, K. V., Hansen, M. H., & Vuust, P. (2023). The effect of music on sleep in hospitalized patients: A systematic review and meta-analysis. Sleep Health. doi:10.1016/j.sleh.2023.03.004

Jespersen, K. V., Otto, M., Kringelbach, M., Van Someren, E., & Vuust, P. (2019). A randomized controlled trial of bedtime music for insomnia disorder. Journal of Sleep Research, 28(4), e12817. doi:10.1111/jsr.12817

Jespersen, K. V., Pando-Naude, V., Koenig, J., Jennum, P., & Vuust, P. (2022). Listening to music for insomnia in adults. Cochrane Database of Systematic Reviews(8). doi:10.1002/14651858.CD010459.pub3.

Jobert, M., Schulz, H., Jähnig, P., Tismer, C., Bes, F., & Escola, H. (1994). A computerized method for detecting episodes of wakefulness during sleep based on the alpha slow-wave index (ASI). Sleep, 17(1), 37–46.

Kakar, E., Venema, E., Jeekel, J., Klimek, M., & van der Jagt, M. (2021). Music intervention for sleep quality in critically ill and surgical patients: a meta-analysis. BMJ Open, 11(5), e042510. doi:10.1136/bmjopen-2020-042510

Kaneshiro, B., Nguyen, D. T., Norcia, A. M., Dmochowski, J. P., & Berger, J. (2020). Natural music evokes correlated EEG responses reflecting temporal structure and beat. Neuroimage, 214, 116559.

Lacaux, C., Strauss, M., Bekinschtein, T. A., & Oudiette, D. (2024). Embracing sleep-onset complexity. Trends in Neurosciences.

Lazic, S. E., & Ogilvie, R. D. (2007). Lack of efficacy of music to improve sleep: A polysomnographic and quantitative EEG analysis. International Journal of Psychophysiology, 63(3), 232–239. doi:10.1016/j.ijpsycho.2006.10.004

Lenc, T., Keller, P. E., Varlet, M., & Nozaradan, S. (2020). Neural and behavioral evidence for frequency-selective context effects in rhythm processing in humans. Cerebral Cortex Communications, 1(1), tgaa037.

Li, J., Ilina, A., Peach, R., Wei, T., Rhodes, E., Jaramillo, V., … Grossman, N. (2025). Falling asleep follows a predictable bifurcation dynamic. Nature Neuroscience, 1–11.

Mayayo, F., Celma-Miralles, A., Keller, P. E., & Toro, J. M. (2026). The role of competing grouping patterns and tonal coherence in neural synchronization to musical meter. Experimental Brain Research, 244(1), 18.

Morin, C. M., LeBlanc, M., Daley, M., Gregoire, J. P., & Mérette, C. (2006). Epidemiology of insomnia: Prevalence, self-help treatments, consultations, and determinants of help-seeking behaviors. Sleep Medicine, 7(2), 123–130. doi:10.1016/j.sleep.2005.08.008

Møller, C., Celma-Miralles, A., Borges, H. B., Stupacher, J., Christensen, C. B., Vuust, P., & Kidmose, P. (2026). Polyrhythms in the Brain: Metrical Priming, Acoustic Balance, and Perceptual Biases. Annals of the New York Academy of Sciences, 1559(1), e70290.

Møller, C., Stupacher, J., Celma-Miralles, A., & Vuust, P. (2021). Beat perception in polyrhythms: Time is structured in binary units. PloS One, 16(8), e0252174.

Nozaradan, S. (2012). Selective neuronal entrainment to the beat and meter embedded in a musical rhythm. The Journal of neuroscience, 32(49), 17572–17581. doi:10.1523/JNEUROSCI.3203-12.2012

Nozaradan, S. (2014). Exploring how musical rhythm entrains brain activity with electroencephalogram frequency-tagging. Philosophical Transactions of the Royal Society B: Biological Sciences, 369(1658), 20130393.

Nozaradan, S., Peretz, I., & Mouraux, A. (2012). Steady-state evoked potentials as an index of multisensory temporal binding. Neuroimage, 60(1), 21–28.

Ollen, J. E. (2006). A criterion-related validity test of selected indicators of musical sophistication using expert ratings. The Ohio State University,

Pallesen, S., Sivertsen, B., Nordhus, I. H., & Bjorvatn, B. (2014). A 10-year trend of insomnia prevalence in the adult Norwegian population. Sleep Medicine, 15(2), 173–179.

Perslev, M., Darkner, S., Kempfner, L., Nikolic, M., Jennum, P. J., & Igel, C. (2021). U-Sleep: resilient high-frequency sleep staging. NPJ digital medicine, 4(1), 72.

Prerau, M. J., Hartnack, K. E., Obregon-Henao, G., Sampson, A., Merlino, M., Gannon, K., … Purdon, P. L. (2014). Tracking the sleep onset process: an empirical model of behavioral and physiological dynamics. PLoS Computational Biology, 10(10), e1003866.

Retter, T. L., Rossion, B., & Schiltz, C. (2021). Harmonic amplitude summation for frequency-tagging analysis. Journal of Cognitive Neuroscience, 33(11), 2372–2393.

Sauvé, S. A., Bolt, E. L., Nozaradan, S., & Zendel, B. R. (2022). Aging effects on neural processing of rhythm and meter. Frontiers in Aging Neuroscience, 14, 848608.

Sifuentes-Ortega, R., Lenc, T., Nozaradan, S., & Peigneux, P. (2022). Partially preserved processing of musical rhythms in REM but not in NREM sleep. Cerebral Cortex, 32(7), 1508–1519.

Sivertsen, B., Harvey, A. G., Vedaa, Ø., Pallesen, S., & Hysing, M. (2026). Sleep Across the Pandemic in Norwegian University and College Students: A National Repeated Cross-Sectional Analysis (2010–2023). Journal of Sleep Research, n/a(n/a), e70312. doi:10.1111/jsr.70312

Strauss, M., Sitt, J. D., Naccache, L., & Raimondo, F. (2022). Predicting the loss of responsiveness when falling asleep in humans. Neuroimage, 251, 119003.

Stupacher, J., Witte, M., Hove, M. J., & Wood, G. (2016). Neural entrainment in drum rhythms with silent breaks: evidence from steady-state evoked and event-related potentials. Journal of Cognitive Neuroscience, 28(12), 1865–1877.

Su, C. P., Lai, H. L., Chang, E. T., Yiin, L. M., Perng, S. J., & Chen, P. W. (2013). A randomized controlled trial of the effects of listening to non-commercial music on quality of nocturnal sleep and relaxation indices in patients in medical intensive care unit. Journal of Advanced Nursing, 69(6), 1377–1389. doi:10.1111/j.1365-2648.2012.06130.x

Šušmáková, K., & Krakovská, A. (2007). Classification of waking, sleep onset and deep sleep by single measures. Sci. Rev, 7, 34–38.

Trahan, T., Durrant, S. J., Müllensiefen, D., & Williamson, V. J. (2018). The music that helps people sleep and the reasons they believe it works: A mixed methods analysis of online survey reports. PloS One, 13(11), e0206531. doi:10.1371/journal.pone.0206531

Trehub, S. E., & Trainor, L. (1998). Singing to infants: Lullabies and play songs. Advances in infancy research, 12, 43–78.

Zhao, N., Lund, H. N., & Jespersen, K. V. (2024). A systematic review and meta-analysis of music interventions to improve sleep in adults with mental health problems. European Psychiatry, 67(1), e62.

