## Supplementary material for "Music facilitates sleep initiation in adults with sleep-onset insomnia: The role of neural synchronization"

### Supplementary materials

Kira Vibe Jespersen*[,1], Alexandre Celma-Miralles*[,1], Peter Vuust[1]

*Shared first-authorship

[1]Center for Music in the Brain, Department of Clinical Medicine, Health, Aarhus University and The Royal Academy of Music Aarhus/Aalborg, Aarhus, Denmark

**Corresponding author:**

Kira Vibe Jespersen,

**Content:**

**Table S1** – The music stimuli of the two playlists including track information, duration, tempo, beat stability and beat-related neural amplitudes.

**Table S2** - Model parameters for the mixed effect model evaluating change in the Delta-Alpha ratio by group.

**Table S3** - Model parameters for the mixed effect model evaluating change in the Percentage of time scored as sleep by group.

**Table S4 -** Model parameters for the mixed effect model evaluating relative Delta power between groups.

**Table S5** - Model parameters for the mixed effect model evaluating relative Theta power between groups.

**Table S6** - Model parameters for the mixed effect model evaluating relative Alpha power between groups.

**Table S7** - Model parameters for the mixed effect model evaluating relative Beta power between groups.

**Table S8** - Model parameters for the mixed effect model evaluating relative Gamma power between groups.

**Table S9** - Evaluation of whether beat-related neural amplitudes were different from zero.

**Figure S1**. - Relative power in percentage across frequency bands.

**Figure S2** - Tapping responses to the beat of the music tracks.

**Figure S3** - Percentage of time spent asleep for each segment in each group.

**Figure S4** - The relationship between beat-related amplitude and sleep initiation.

**Table S1.** The music stimuli of the two playlists including track information, duration, tempo, beat stability and beat-related neural amplitudes. Mean values (±SD).

| No | Title | Duration (minutes) | Tempo (Hz) | Tempo (BPM) | Beat stability (tapping)[1] | 1st session EEG[2] amplitudes (µV) | 2nd session EEG amplitudes (µV) | Significant peak |
|---|---|---|---|---|---|---|---|---|
| **Minimalistic Playlist** | | | | | | | | |
| 1 | Fyrsta[a] | 4.16 | 1.0783 | 65 | 0,097 (±0.017) | -0,003 (±0.024) | 0,007 (±0.001) | No |
| 2 | This Place Is a Shelter[a] | 2.18 | 0.7392 | 44 | 0,283 (±0.104) | -0,004 (±0.059) | 0,008 (±0.039) | No |
| 3 | Ágúst[a] | 3.06 | 1.0835 | 65 | 0,843 (±0.175) | 0,087 (±0.067) | 0,086 (±0.086) | Yes |
| 4 | Nyepi[b] | 4.14 | 1.0152 | 61 | 0,155 (±0.045) | 0,004 (±0.050) | -0,003 (±0.027) | No |
| 5 | Saman[b] | 2.11 | 0.9101 | 55 | 0,467 (±0.153) | 0,004 (±0.043) | -0,028 (±0.062) | No |
| 6 | Tomorrow's Song[a] | 3.07 | 1.1729 | 70 | 0,388 (±0.124) | 0,013 (±0.035) | 0,013 (±0.032) | No |
| 7 | You[c] | 3.06 | 0.9416 | 57 | 0,383 (±0.116) | -0,014 (±0.070) | -0,024 (±0.029) | No |
| 8 | Do[c] | 3.04 | 0.6787 | 41 | 0,455 (±0.221) | -0,011 (±0.048) | -0,038 (±0.066) | No |
| 9 | Re[c] | 3.50 | 1.1893 | 71 | 0,135 (±0.047) | -0,008 (±0.056) | -0,013 (±0.060) | No |
| 10 | Mi[c] | 3.59 | 0.7205 | 43 | 0,099 (±0.048) | 0,007 (±0.063) | -0,014 (±0.052) | No |
| **New Age Playlist** | | | | | | | | |
| 1 | The North[d] | 2.36 | 0.8679 | 52 | 0,667 (±0.173) | 0,04 (±0.059) | 0,03 (±0.037) | Yes |
| 2 | The North[d] | 3.24 | 0.8679 | 52 | 0,666 (±0.154) | 0,042 (±0.043) | 0,014 (±0.059) | Yes |
| 3 | The North[d] | 4.28 | 0.8679 | 52 | 0,679 (±0.159) | 0,032 (±0.003) | 0,026 (±0.018) | Yes |
| 4 | The North[d] | 3.50 | 0.8679 | 52 | 0,663 (±0.168) | 0,025 (±0.044) | 0,042 (±0.055) | Yes |
| 5 | The North[d] | 2.39 | 1.1667 | 70 | 0,714 (±0.213) | 0,003 (±0.063) | 0,053 (±0.102) | Yes |
| 6 | The Cave[e] | 1.14 | 0.8679 | 52 | 0,914 (±0.068) | 0,054 (±0.080) | 0,007 (±0.096) | Yes |
| 7 | The Cave[e] | 4.32 | 0.8679 | 52 | 0,644 (±0.237) | 0,044 (±0.067) | 0,047 (±0.095) | Yes |
| 8 | The Cave[e] | 2.42 | 0.8679 | 52 | 0,841 (±0.149) | 0,016 (±0.122) | 0,023 (±0.040) | No |
| 9 | The Cave[e] | 4.41 | 0.8679 | 52 | 0,791 (±0.058) | 0,028 (±0.054) | -0,022 (±0.066) | No |
| 10 | The Cave[e] | 4.30 | 0.8679 | 52 | 0,709 (±0.191) | 0,039 (±0.045) | 0,023 (±0.039) | Yes |

Notes

[a]Ólafur Arnalds, Living room songs, Erased Tapes Records, 2011.

[b]Ólafur Arnalds, re:member, Mercury KX, 2018.

[c]Nils Frahm, Screws, Erased Tapes Records, 2012.

[d]Niels Eje, Musicure 4 – Northern Light, Gefion Records, 2004.

[e]Niels Eje, Musicure 6 – Waves, Gefion Records, 2006.

[1] Beat stability was determined via finger tapping responses. Tapping consistency at each tempo was obtained from the mean of all participants' tapping vector length, for which 0 indicates 'random tapping' and 1 indicates 'isochronous tapping'', and its reliability tested with Rayleigh tests.

[2] Beat-related neural amplitudes derived from frequency-tagging the EEG.

| **Table S2.** Model parameters for the mixed effect model evaluating change in the Delta-Alpha ratio by group. | | | | |
|---|---|---|---|---|
| Model equation: LOGrel_ratio_deltaalpha ~ Session + Group * poly(track, 2) + (Session \| Participant) + (1 \| track) | | | | |
| Fixed Effects | | | | |
| | Estimate | SE | t | p |
| Intercept | 0.0009 | 0.1596 | 0.006 | 0.995 |
| Music group | 0.2234 | 0.2238 | 0.998 | 0.323 |
| Session | 0.0208 | 0.0962 | 0.216 | 0.830 |
| **poly(track, 2)1** | **6.018** | **1.280** | **4.702** | **0.001** |
| **poly(track, 2)2** | **-5.130** | **1.280** | **-4.008** | **0.002** |
| **Music group × poly(track, 2)1** | **2.976** | **1.248** | **2.384** | **0.017** |
| Music group × poly(track, 2)2 | 2.018 | 1.248 | 1.616 | 0.106 |
| Random Effects | | | | |
| | | | Variance | S.D. |
| Participant (Intercept) | | | 0.6756 | 0.8220 |
| Session | | | 0.3174 | 0.5634 |
| Track (Intercept) | | | 0.0102 | 0.1010 |
| Residual | | | 0.3832 | 0.6191 |

*Notes:*
Significant effects are highlighted in bold.
The model was estimated using restricted maximum likelihood (REML).
Session reflects the time of the measurement, i.e. baseline session or follow-up session after 4-weeks of music listening.
Music reflects the Group parameter, i.e. music or silence.
Poly(track, 2) reflects the Time parameter evaluating both a linear (1) and quadratic fit (2).
Track reflects the 3-minute segments of the 30-minute recording.

**Table S3.** Model parameters for the mixed effect model evaluating change in the Percentage of time scored as sleep by group.

Model equation: SleepPercentage ~ Session + Group * poly(track, 2) + (Session | Participant) + (1 | track)

Fixed Effects

| | Estimate | SE | t | p |
|---|---|---|---|---|
| Intercept | 16.1943 | 4.5102 | 3.591 | <0.001 |
| Music group | 4.4528 | 5.6819 | 0.784 | 0.437 |
| Session | -0.9836 | 4.4825 | -0.219 | 0.827 |
| **poly(track, 2)1** | **158.0996** | **44.1329** | **3.582** | **0.004** |
| **poly(track, 2)2** | **-186.2889** | **44.1329** | **-4.221** | **0.001** |
| **Music group × poly(track, 2)1** | **121.9907** | **48.3938** | **2.521** | **0.012** |
| Music group × poly(track, 2)2 | 29.5207 | 48.3938 | 0.610 | 0.542 |

Random Effects

| | Variance | S.D. |
|---|---|---|
| Participant (Intercept) | 550.647 | 23.466 |
| Session | 796.414 | 28.221 |
| Track (Intercept) | 9.853 | 3.139 |
| Residual | 575.948 | 23.999 |

*Notes:*
Significant effects are highlighted in bold.
The model was estimated using restricted maximum likelihood (REML).
Session reflects the time of the measurement, i.e. baseline session or follow-up session after 4-weeks of music listening.
Music reflects the Group parameter, i.e. music or silence.
Poly(track, 2) reflects the Time parameter evaluating both a linear (1) and quadratic fit (2).
Track reflects the 3-minute segments of the 30-minute recording.

**Table S4.** Model parameters for the mixed effect model evaluating relative Delta power between groups.

Model equation: rel_delta ~ Session + MusicGroup * poly(track, 2) + (Session|Participant) + (1 | track)

| Fixed Effects | | | | |
|---|---|---|---|---|
| | Estimate | SE | t | p |
| Intercept | 31.6667 | 2.3520 | 13.464 | <0.001 |
| Music group | -2.8573 | 2.9685 | -0.963 | 0.340 |
| Session | 0.4026 | 1.2531 | 0.321 | 0.750 |
| **poly(track, 2)1** | **115.6339** | **15.5908** | **7.417** | **<0.001** |
| poly(track, 2)2 | -24.7576 | 15.5908 | -1.588 | 0.119 |
| Music group × poly(track, 2)1 | -40.7977 | 22.0830 | -1.847 | 0.079 |
| Music group × poly(track, 2)2 | -33.2442 | 22.0830 | -1.505 | 0.147 |

| Random Effects | | |
|---|---|---|
| | Variance | S.D. |
| Participant (Intercept) | 127.06 | 11.27 |
| Session | 49.15 | 7.01 |
| Track (Intercept) | 0.98 | 0.99 |
| Residual | 85.77 | 9.26 |

*Notes:*
Significant effects are highlighted in bold.
The model was estimated using restricted maximum likelihood (REML).
Session reflects the time of the measurement, i.e. baseline session or follow-up session after 4-weeks of music listening.
Music reflects the Group parameter, i.e. music or silence.
Poly(track, 2) reflects the Time parameter evaluating both a linear (1) and quadratic fit (2).
Track reflects the 3-minute segments of the 30-minute recording.

| **Table S5.** Model parameters for the mixed effect model evaluating relative Theta power between groups. | | | | |
|---|---|---|---|---|
| Model equation: rel_theta ~ Session + MusicGroup * poly(track, 2) + (Session\|Participant) + (1 \| track) | | | | |
| Fixed Effects | | | | |
| | Estimate | SE | t | p |
| Intercept | 19.0555 | 1.0470 | 18.201 | <0.001 |
| Music group | -0.5770 | 1.2773 | -0.452 | 0.653 |
| Session | -0.6979 | 0.6005 | -1.162 | 0.252 |
| poly(track, 2)1 | 8.6394 | 8.0843 | 1.069 | 0.293 |
| **poly(track, 2)2** | **-33.9589** | **8.0843** | **-4.201** | **<0.001** |
| Music group × poly(track, 2)1 | -0.1162 | 12.4827 | -0.009 | 0.993 |
| Music group × poly(track, 2)2 | -0.7632 | 12.4827 | -0.061 | 0.952 |
| Random Effects | | | | |
| | | | Variance | S.D. |
| Participant (Intercept) | | | 26.8351 | 5.1803 |
| Session | | | 12.4006 | 3.5215 |
| Track (Intercept) | | | 0.6934 | 0.8327 |
| Residual | | | 14.2529 | 3.7753 |

*Notes:*
Significant effects are highlighted in bold.
The model was estimated using restricted maximum likelihood (REML).
Session reflects the time of the measurement, i.e. baseline session or follow-up session after 4-weeks of music listening.
Music reflects the Group parameter, i.e. music or silence.
Poly(track, 2) reflects the Time parameter evaluating both a linear (1) and quadratic fit (2).
Track reflects the 3-minute segments of the 30-minute recording.

**Table S6.** Model parameters for the mixed effect model evaluating relative Alpha power between groups.

Model equation: rel_alpha ~ Session + MusicGroup * poly(track, 2) + (Session|Participant) + (1 | track)

Fixed Effects

| | Estimate | SE | t | p |
|---|---|---|---|---|
| Intercept | 27.2771 | 2.6444 | 10.315 | <0.001 |
| Music group | 1.9819 | 3.4805 | 0.569 | 0.571 |
| Session | 0.6497 | 1.4852 | 0.437 | 0.664 |
| **poly(track, 2)1** | **-142.1467** | **17.0376** | **-8.343** | **<0.001** |
| **poly(track, 2)2** | **54.9197** | **17.0376** | **3.223** | **0.002** |
| Music group × poly(track, 2)1 | 46.1108 | 25.2672 | 1.825 | 0.081 |
| Music group × poly(track, 2)2 | 24.3390 | 25.2672 | 0.963 | 0.346 |

Random Effects

| | Variance | S.D. |
|---|---|---|
| Participant (Intercept) | 144.169 | 12.007 |
| Session | 78.882 | 8.882 |
| Track (Intercept) | 2.149 | 1.466 |
| Residual | 82.402 | 9.078 |

*Notes:*
Significant effects are highlighted in bold.
The model was estimated using restricted maximum likelihood (REML).
Session reflects the time of the measurement, i.e. baseline session or follow-up session after 4-weeks of music listening.
Music reflects the Group parameter, i.e. music or silence.
Poly(track, 2) reflects the Time parameter evaluating both a linear (1) and quadratic fit (2).
Track reflects the 3-minute segments of the 30-minute recording.

| **Table S7.** Model parameters for the mixed effect model evaluating relative Beta power between groups. | | | | |
|---|---|---|---|---|
| Model equation: rel_beta ~ Session + MusicGroup * poly(track, 2) + (Session\|Participant) + (1 \| track) | | | | |
| Fixed Effects | | | | |
| | Estimate | SE | t | p |
| Intercept | 18.7464 | 1.6519 | 11.349 | <0.001 |
| Music group | 2.1105 | 2.1949 | 0.962 | 0.341 |
| Session | -0.3381 | 0.4777 | -0.708 | 0.483 |
| poly(track, 2)1 | 2.8778 | 6.4999 | 0.443 | 0.658 |
| poly(track, 2)2 | -1.1288 | 6.4999 | -0.174 | 0.862 |
| Music group × poly(track, 2)1 | 0.6791 | 8.6564 | 0.078 | 0.937 |
| Music group × poly(track, 2)2 | 5.3502 | 8.6564 | 0.618 | 0.537 |
| Random Effects | | | | |
| | | | Variance | S.D. |
| Participant (Intercept) | | | 63.567 | 7.973 |
| Session | | | 5.756 | 2.399 |
| Track (Intercept) | | | 0.000 | 0.000 |
| Residual | | | 18.428 | 4.293 |

*Notes:*
Significant effects are highlighted in bold.
The model was estimated using restricted maximum likelihood (REML).
Session reflects the time of the measurement, i.e. baseline session or follow-up session after 4-weeks of music listening.
Music reflects the Group parameter, i.e. music or silence.
Poly(track, 2) reflects the Time parameter evaluating both a linear (1) and quadratic fit (2).
Track reflects the 3-minute segments of the 30-minute recording.

| **Table S8.** Model parameters for the mixed effect model evaluating relative Gamma power between groups. | | | | |
|---|---|---|---|---|
| Model equation: rel_gamma ~ Session + MusicGroup * poly(track, 2) + (Session\|Participant) + (1 \| track) | | | | |
| Fixed Effects | | | | |
| | Estimate | SE | t | p |
| Intercept | 3.2159 | 0.4407 | 7.298 | <0.001 |
| Music group | -0.4380 | 0.5547 | -0.790 | 0.433 |
| Session | -0.1981 | 0.2897 | -0.684 | 0.498 |
| **poly(track, 2)1** | **14.6351** | **3.5939** | **4.072** | **<0.001** |
| poly(track, 2)2 | 4.3597 | 3.5939 | 1.213 | 0.225 |
| Music group × poly(track, 2)1 | -5.4953 | 4.7862 | -1.148 | 0.251 |
| Music group × poly(track, 2)2 | 4.8861 | 4.7862 | 1.021 | 0.308 |
| Random Effects | | | | |
| | | | Variance | S.D. |
| Participant (Intercept) | | | 3.940 | 1.985 |
| Session | | | 2.511 | 1.584 |
| Track (Intercept) | | | 0.000 | 0.000 |
| Residual | | | 5.634 | 2.374 |

*Notes:*
Significant effects are highlighted in bold.
The model was estimated using restricted maximum likelihood (REML).
Session reflects the time of the measurement, i.e. baseline session or follow-up session after 4-weeks of music listening.
Music reflects the Group parameter, i.e. music or silence.
Poly(track, 2) reflects the Time parameter evaluating both a linear (1) and quadratic fit (2).
Track reflects the 3-minute segments of the 30-minute recording.

**Table S9.** Evaluation of whether beat-related neural amplitudes were different from zero

Model equation: BeatAmplitude ~ 0 + Track + (Session | Participant)

Fixed Effects

| | Estimate | SE | t | p |
|---|---|---|---|---|
| Track 1 Minimalistic | 0.0007 | 0.0136 | 0.054 | 0.957 |
| Track 2 Minimalistic | 0.0005 | 0.0136 | 0.037 | 0.971 |
| **Track 3 Minimalistic** | **0.0867** | **0.0136** | **6.365** | **<0.001** |
| Track 4 Minimalistic | 0.0012 | 0.0136 | 0.091 | 0.927 |
| Track 5 Minimalistic | -0.0077 | 0.0136 | -0.563 | 0.574 |
| Track 6 Minimalistic | 0.0134 | 0.0136 | 0.985 | 0.325 |
| Track 7 Minimalistic | -0.018 | 0.0136 | -1.322 | 0.187 |
| Track 8 Minimalistic | -0.021 | 0.0136 | -1.541 | 0.124 |
| Track 9 Minimalistic | -0.0096 | 0.0136 | -0.708 | 0.479 |
| Track 10 Minimalistic | -0.0008 | 0.0136 | -0.060 | 0.952 |
| **Track 1 New Age** | **0.0346** | **0.0127** | **2.734** | **0.007** |
| **Track 2 New Age** | **0.0277** | **0.0127** | **2.190** | **0.029** |
| **Track 3 New Age** | **0.0288** | **0.0127** | **2.279** | **0.023** |
| **Track 4 New Age** | **0.0338** | **0.0127** | **2.672** | **0.023** |
| **Track 5 New Age** | **0.0283** | **0.0127** | **2.233** | **0.026** |
| **Track 6 New Age** | **0.0304** | **0.0127** | **2.403** | **0.017** |
| **Track 7 New Age** | **0.0455** | **0.0127** | **3.598** | **<0.001** |
| Track 8 New Age | 0.0195 | 0.0127 | 1.537 | 0.125 |
| Track 9 New Age | 0.0029 | 0.0127 | 0.229 | 0.819 |
| **Track 10 New Age** | **0.0310** | **0.0127** | **2.448** | **0.015** |

Random Effects

| | Variance | S.D. |
|---|---|---|
| Participant (Intercept) | <0.0000 | <0.0000 |
| Session | <0.0000 | <0.0000 |
| Residual | 0.0035 | 0.0594 |

*Notes:*

Significant effects are highlighted in bold.

The model was estimated using restricted maximum likelihood (REML).

Session reflects the time of the measurement, i.e. baseline session or follow-up session after 4-weeks of music listening.

Track reflects the music track sleep playlist numbered 1-10 (Minimalistic) and 11-20 (New Age)

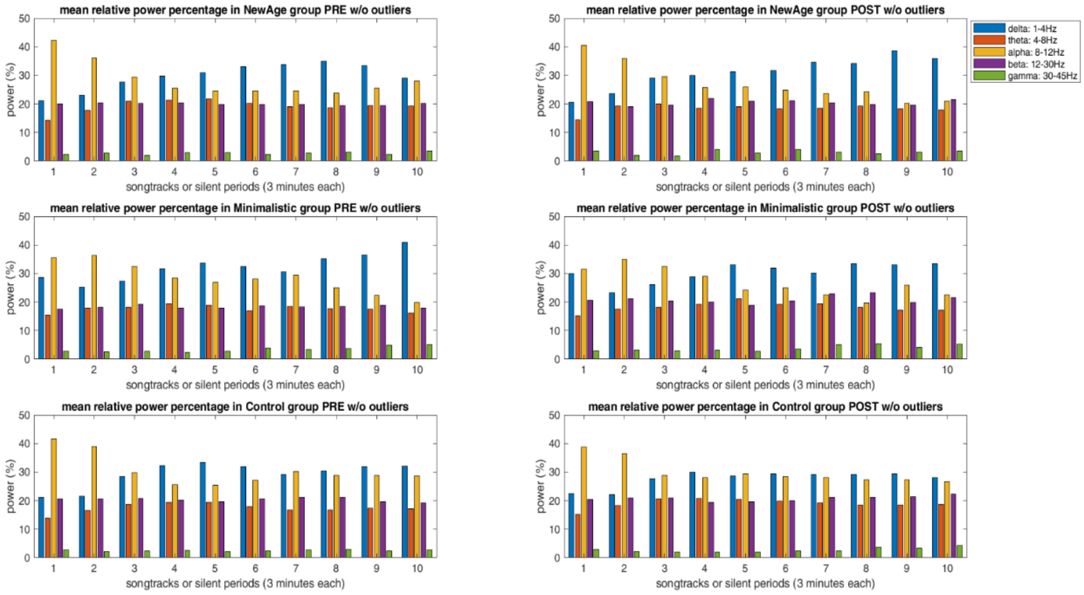

**Figure S1. Power Spectral Density measures from EEG recordings.**

Relative power in percentage across frequency bands (i.e., delta, theta, alpha, beta, and gamma) in the music listening group (top and middle row) and the silently resting group (bottom row) in the first (left) and second (right) EEG session. Note how alpha activity (yellow bars) decreases over the course of the ten 3-minute tracks, while delta activity (blue bars) increases over time.

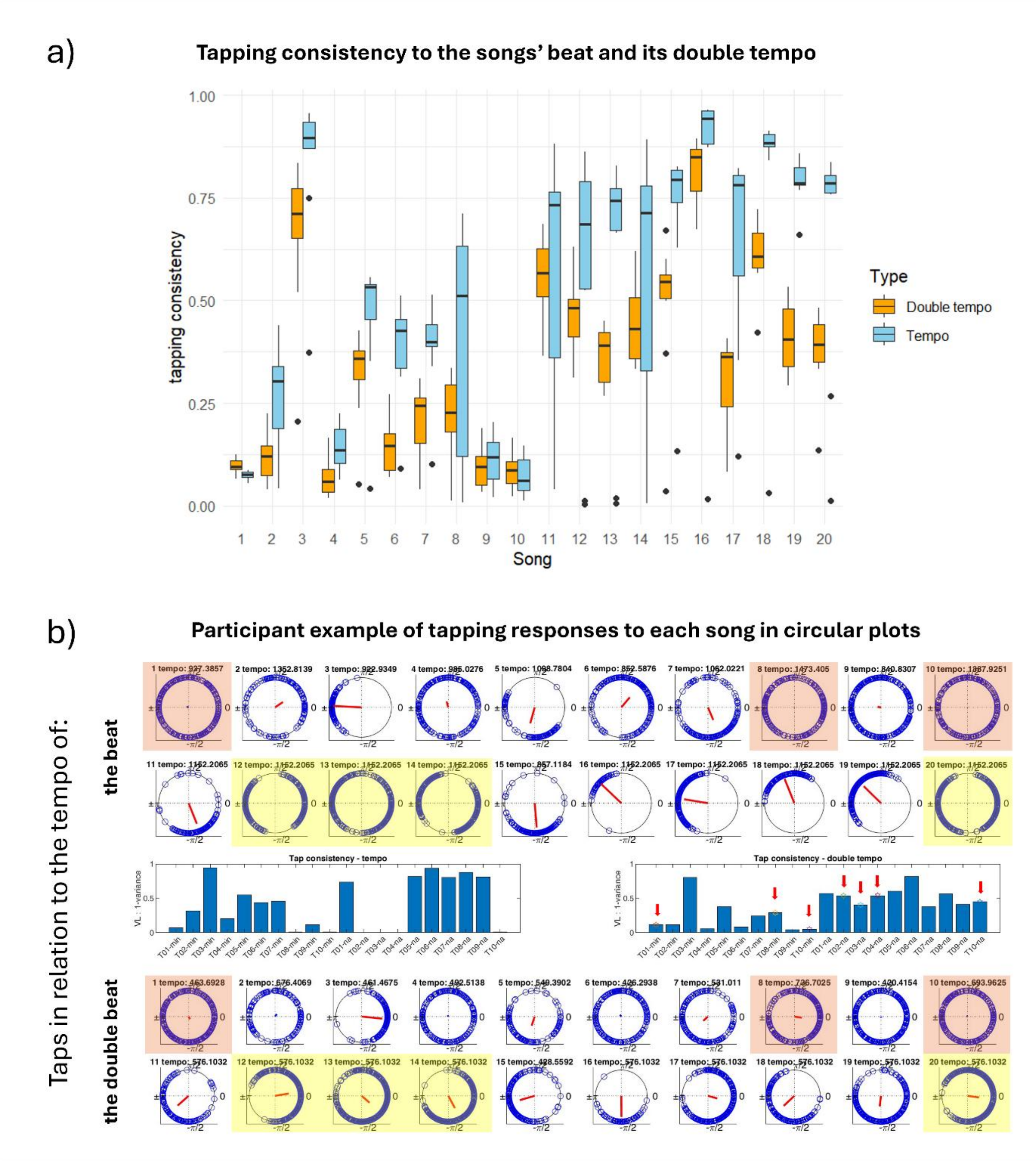

**Figure S2. Tapping responses to the beat of the music tracks.**

(a) Tapping consistency for each song across participants tapping to the perceived beat with respect to the tempo (blue) or double tempo (orange). Higher scores on the y-axis indicate more stable tapping at the respective beat. On the x-axis music tracks 1-10 are the Minimalistic playlist and tracks 11-20 are the New Age Playlist. (b) An example of the tapping of one participant including circular representations of the tapping data with blue circles representing each tap and the red line indicating the mean vector length, i.e. the stability of the taps calculated as 1-variance. In some tracks, the participant tapped to the double beat (circular plots highlighted in salmon for tracks 1-10, the Minimalistic playlist, and in yellow for the tracks 11-20, the New Age playlist). In these cases, we took the vector length for the double beat to calculate beat stability for the given track (marked with arrows in the example bar plot), since it revealed less variance (i.e. longer red vector) and therefore a better approximation to beat consistency.

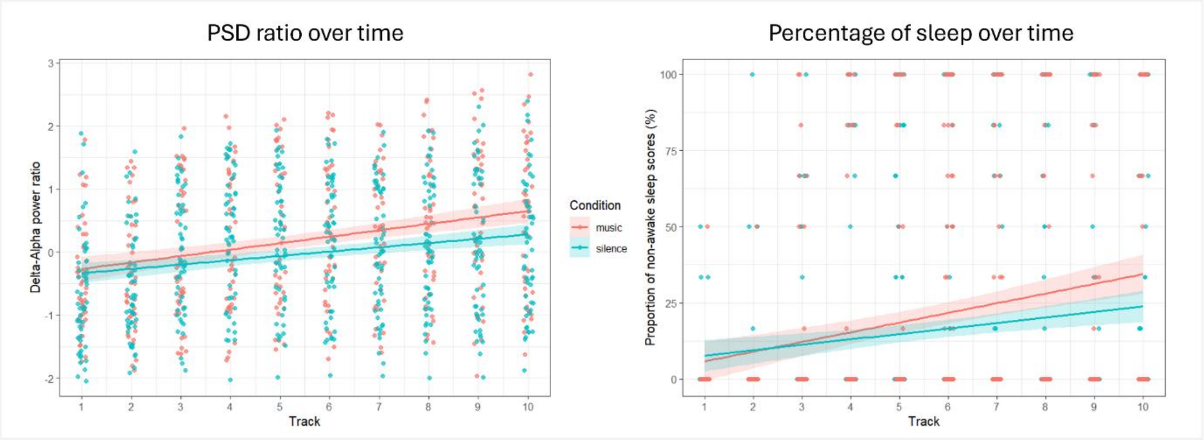

**Figure S3. Percentage of time spent asleep for each segment in each group.**
To further investigate sleep onset, we did sleep scoring of the 30-minute EEG recording using the U-sleep automated sleep scoring algorithm (see Methods section). For each 3-minute segment we calculated the percentage of time scored as sleep. Because this study focused on sleep onset during a 30-minute rest, we mainly found N1 and N2 sleep stages, and we distinguished between wakefulness and sleep rather than focusing on the sleep stages. The results showed the same pattern as for the Delta-Alpha ratio. There was a main effect of Time (Track) and a significant Group x Time interaction with the Music group (salmon) showing a higher increase in sleep percentage compared to the Silence group (turquoise). For further details, see Supplementary Table S3.

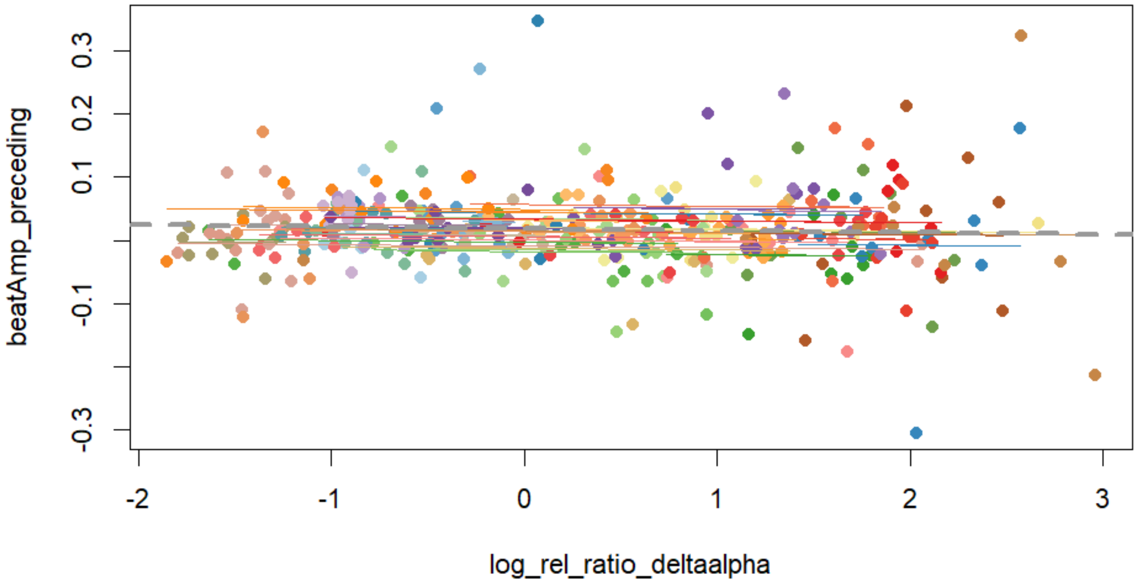

**Figure S4 – The relationship between precedent beat-related amplitude and sleep initiation.** Repeated measures correlations showed no relation between the beat-related amplitude of one track and degree of wakefulness (delta-alpha ratio) in the following track (r = -0.02, 95% CI: -0.13 to 0.08, *p* = 0.6776). As such, our results do not support the hypothesis that sleep initiation is facilitated by neural synchronization to the beat of the preceding music track.
